# Langerhans Cells Drive Tfh and B Cell Responses Independent of Canonical Cytokine Signals

**DOI:** 10.1101/2025.01.10.632426

**Authors:** Aurélie Bouteau, Zhen Qin, Sandra Zurawski, Gerard Zurawski, Botond Z. Igyártó

## Abstract

Dendritic cells (DCs) are key regulators of adaptive immunity, guiding T helper (Th) cell differentiation through antigen presentation, co-stimulation, and cytokine production. However, in steady-state conditions, certain DC subsets, such as Langerhans cells (LCs), induce T follicular helper (Tfh) cells and B cell responses without inflammatory stimuli. Using multiple mouse models and *in vitro* systems, we investigated the mechanisms underlying steady-state LC-induced adaptive immune responses. We found that LCs drive germinal center Tfh and B cell differentiation and antibody production independently of interleukin-6 (IL-6), type-I interferons, and ICOS ligand (ICOS-L) signaling, which are critical in inflammatory settings. Instead, these responses relied on CD80/CD86-mediated co-stimulation. Our findings challenge the conventional three-signal paradigm by demonstrating that cytokine signaling is dispensable for LC-mediated Tfh and B cell responses in steady-state. These insights provide a framework for understanding homeostatic immunity and the immune system’s role in maintaining tolerance or developing autoimmunity under non-inflammatory conditions.

**SUMMARY STATEMENT:** Langerhans cells (LCs) drive germinal center Tfh and B cell responses in steady-state conditions independently of IL-6, type-I interferons, and ICOS ligand, challenging the established cytokine-centric model of T cell differentiation.

## INTRODUCTION

Dendritic cells (DCs) are critical in training and educating naïve T cells and their differentiation into specific T helper subsets (Merad et al., 2013; Yin et al., 2021). Generally, it is widely accepted that the DCs provide three signals to naïve T cells in the form of cognate peptide/MHC, membrane-bound co-stimulation, and soluble cytokines. Out of these, the cytokines, as third signals (Curtsinger et al., 1999), are regarded as key components in T helper cell differentiation into T helper subsets, such as Th1, Th2, Th17, Tfh cells, and others (Yin et al., 2021; Hilligan and Ronchese, 2020). These distinct Th subsets are thought to be induced by the DC-derived polarizing cytokines specific for each Th subset. The Th polarizing cytokines are induced by exposure to various inflammatory stimuli sensed by DC through distinct pathogen-associated molecular pattern receptors (Janeway and Medzhitov, 2002). This oversimplified model of Th differentiation provides a plausible explanation for inflammatory settings, but it is challenging to apply in a broader sense. For example, antigen targeting to different mouse DC subsets in steady-state in the apparent lack of adjuvant and other inflammatory signals induces Tfh cells and antibody responses (Yao et al., 2015; Bouteau et al., 2019; Caminschi and Shortman, 2012; Kato et al., 2015; Corbett et al., 2005; Li et al., 2015; Lahoud et al., 2011; Kato et al., 2020; Caminschi et al., 2008; Kervevan et al., 2021). Furthermore, anti-commensal responses, termed homeostatic immunity, happen regularly in the absence of overt inflammation (Belkaid and Harrison, 2017), justifying the need to understand better the induction mechanism of adaptive immune responses in this non-inflammatory context.

Mouse LCs with monocytic origin but DC functions (Ginhoux et al., 2006; Hoeffel et al., 2012; Doebel et al., 2017) drive Tfh cells and germinal center (GC)-dependent protective antibody responses in steady-state (Yao et al., 2015; Bouteau et al., 2019). They do this irrespective of the nature of the receptor targeted and without signs of activation and maturation (Yao et al., 2015; Bouteau et al., 2019). Similarly, primary human LCs and CD34^+^ cord-blood-derived LCs, unlike monocyte-derived DCs, also support Tfh differentiation and B cell responses *in vitro* without adjuvants upon antigen delivery through Langerin (Kervevan et al., 2021; Bouteau et al., 2019). However, the mechanism by which LCs support adaptive immune responses in steady-state remains elusive. Therefore, here, we set out to define the mechanism. We found that LCs induced Tfh and B cell responses independently from IL-6, type-I interferon, and ICOS-L, which were previously reported to play critical roles in inflammatory settings in driving Tfh cells and antibody responses (Crotty, 2014; Krishnaswamy et al., 2018; Crotty, 2019). LC-induced responses, however, were dependent on CD80/CD86 co-stimulation.

## RESULTS AND DISCUSSION

### LCs, unlike cDC1s, induce GC-Tfh cells and antibody responses in steady-state

We have previously shown using two mouse models that steady-state antigen targeting to LCs, but not cDC1, leads to GC-Tfh formation and antibody responses (Bouteau et al., 2019). In mouse skin, Langerin expression is confined to LCs and CD103^+^/XCR-1^+^ cDC1s (Kaplan, 2017), and thus, we used two mouse models, huLangerin and Batf3^−/−^ mice, to permit targeting antigen to either LCs or cDC1s. The huLangerin mice express human Langerin specifically in LCs (Bobr et al., 2010), allowing antigen targeting to LCs in the presence of cDC1s using anti-<u>human</u> Langerin (Yao et al., 2015; Bouteau et al., 2019; Igyártó et al., 2011). The Batf3^−/−^ mice lack the migratory Langerin-expressing cDC1s. Therefore, the remaining LCs can be specifically targeted using anti-<u>mouse</u> Langerin (Bouteau et al., 2019; Yao et al., 2015). To further strengthen our previous findings and increase rigor, here, we expanded our toolset to include two other mouse models that were more recently generated by the Murphy lab, XCR1-Cre and IRF8^32Δ^, both specifically affecting cDC1s (Durai et al., 2019; Ferris et al., 2020). We bred the XCR-1-Cre to “STOP”-DTA mice to selectively eliminate all the cDC1s (**Fig. 1 A**). The huLangerin-DTA (huL-DTA) mice that lack LCs (Kaplan et al., 2005) were used as controls for cDC1s targeting that does not induce GC-Tfh cells and antibody responses in steady-state (Yao et al., 2015; Bouteau et al., 2019). The mice were then adoptively transferred with CD4 TEα cells and injected with 1 μg of anti-muLangerin-Eα a day later. Four and fourteen days later, the antigen-specific CD4^+^ T cell and B cell responses were characterized using flow cytometry and ELISA, respectively, as described previously (Yao et al., 2015; Bouteau et al., 2019). We found that LCs targeted in XCR-1-Cre-DTA and IRF8^32Δ^ mice, like Batf3^−/−^ mice, induced GC-Tfh cells, in sharp contrast to cDC1 targeting in huL-DTA mice (**Fig. 1 B**). The B cell responses mounted in these two new models followed a largely similar trajectory to Batf3^−/−^ mice but were slightly less pronounced (**Fig. 1 C**). Interestingly, the IRF8^32Δ^ mice did not produce significant levels of antibodies, unlike XCR1-Cre-DTA and Batf3^−/−^ mice (**Fig. 1 D**). This could indicate that this IRF8^32Δ^ deletion might affect GC and plasma cell responses, as has been reported for full IRF8^−/−^ mice (Carotta et al., 2014; Wang et al., 2019). Thus, these data further support our previous observation that LCs, unlike cDC1s, can induce GC-Tfh cells and antibody responses in steady-state, irrespective of the mouse model used.

**Figure 1.**
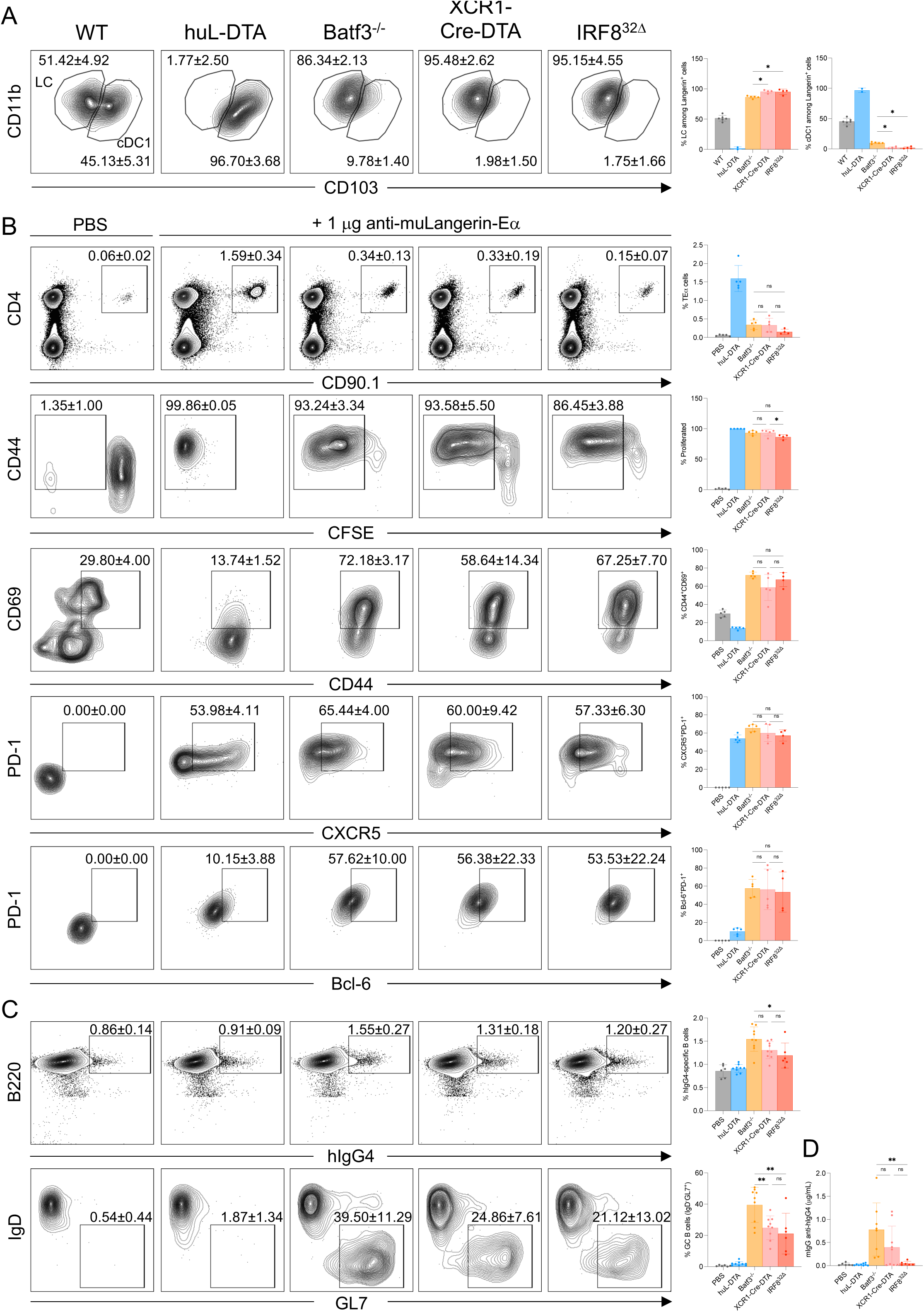
LCs, unlike cDC1s, induce GC-Tfh cells and antibody responses in steady-state. (**A**) LCs and cDC1s were quantified in the indicated mouse strains. Representative flow plots and summary graphs. Upstream gate: live/singlets, MHC-II^hi^, CD11c^+^, Langerin^+^. Data was pooled from two independent experiments. Each dot represents a separate mouse. (**B**) The indicated mice were transferred with TEα cells and then immunized with 1 μg of anti-muLangerin-hIgG4-Eα or vehicle (PBS) the next day. Ag-specific TEα cell responses were assessed 4 days later by flow cytometry. Representative flow plots with corresponding summary graphs are shown. Upstream gate: live/singlets. TEα cells were identified as CD4^+^CD90.1^+^. For the subsequent plots, % of total TEα is shown. Data was pooled from two independent experiments. Each dot represents a separate mouse. (**C**) The mice were immunized with 1 μg of anti-huLangerin-hIgG4-Eα or vehicle (PBS), and fourteen days later, Ag-specific B cell responses were assessed by flow cytometry and (**D**) ELISA. Representative flow plots with corresponding summary graphs are shown. Upstream gate: live/singlets/dump. hIgG4-specific B cells were identified as B220^+^hIgG4^+^. For the subsequent plots, % of total hIgG4 is shown. Data was pooled from three independent experiments. Each dot represents a separate mouse. *p<0.05, **p<0.005, ns=not significant.

### Type I interferon is not required for the induction of adaptive immune responses by steady-state LCs

It is unknown how adaptive immune responses, including T follicular helper cells and antibody responses, are induced by LCs and some DC subsets in steady-state. Intact type-I interferon signaling in DCs is needed to effectively induce Tfh cells in inflammatory settings through IL-6 up-regulation (Cucak et al., 2009). Targeted delivery of IFNα to cDC1 through Langerin enables these cells to support the differentiation of GC-Tfh cells and GC-dependent antibody responses (Bouteau et al., 2019). Based on these data, we hypothesized that steady-state levels of type-I interferon signaling in LCs might play a role in inducing Tfh cells and B cell responses in steady-state. To test this hypothesis, we bred the huLangerinCre mice to IFNαR1^f/f^ mice to delete IFNαR1 from LCs (**Fig. 2 A**). The genotypes of the resulting mice were determined using standard PCR, while the selective deletion of IFNαR1 protein on LCs was confirmed using flow cytometry (**Fig. S1 A**). Cre-positive and cre-negative mice were then adoptively transferred with CD4^+^ TEα cells and, a day later, were injected with 1 μg of anti-huLangerin-Eα. Four and fourteen days later, the antigen-specific CD4^+^ T cell and B cell responses were characterized using flow cytometry and ELISA. The anti-huIgG IgG levels were determined using ELISA on serum samples harvested on day 14 (**Fig. 2 A**). We found no major differences regarding antigen-specific CD4^+^ T cell and B cell expansions and phenotype. The percent of Tfh cells slightly decreased but this was not reflected in the B cell responses. (**Fig. 2, B and C**). The anti-huIgG4 IgG levels were also unaffected without type-I interferon signaling in LCs (**Fig. 2 D**). These data, therefore, point to the lack of a critical role of type-I interferon signaling in LCs in the steady-state induction of Tfh cells and antibody responses.

**Figure 2.**
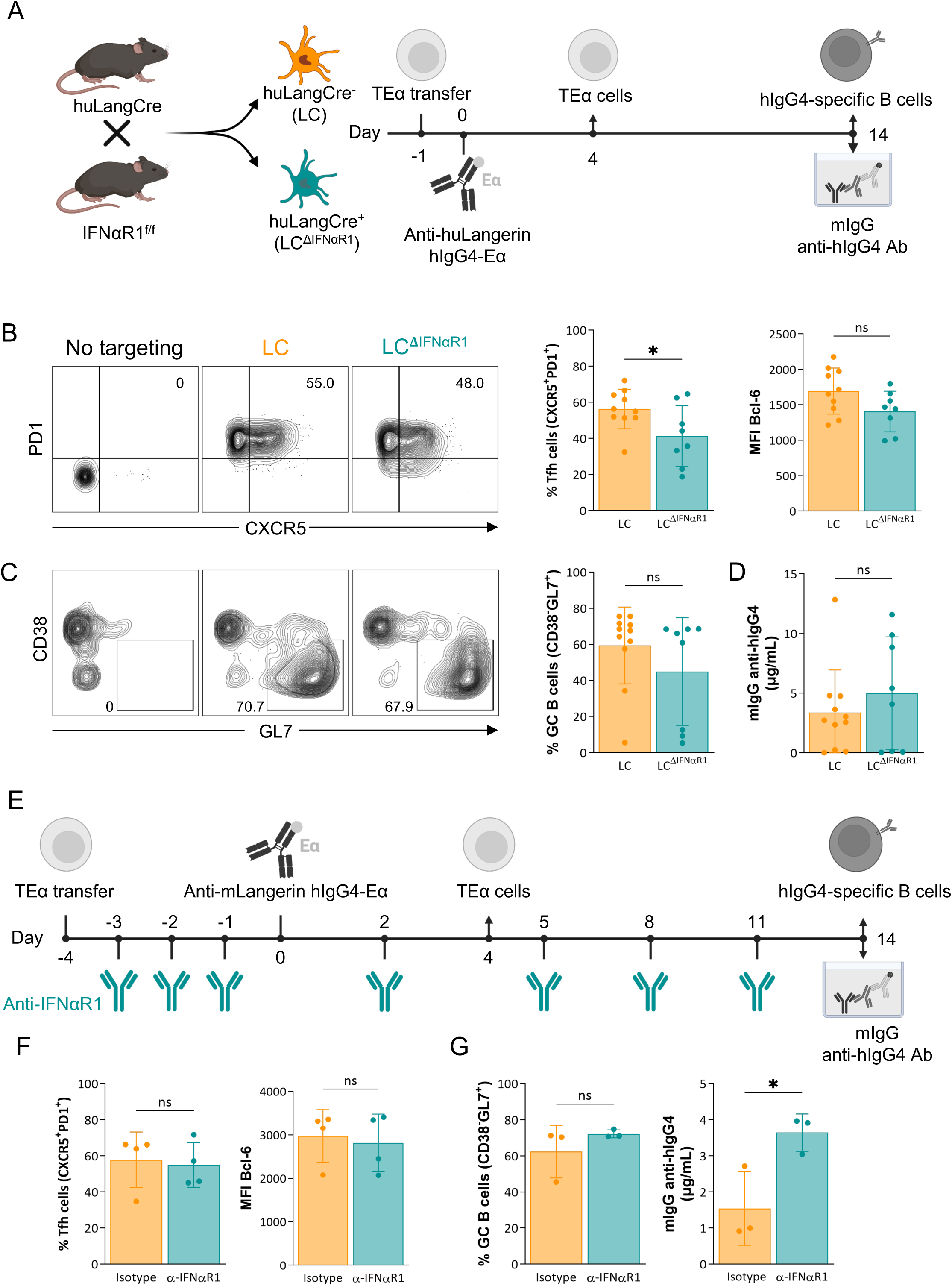
**Type I interferon is not required for the induction of adaptive immune responses by steady-state LCs**. (**A**) huLangCre-IFNαR1^f/f^ mice were generated as depicted, which allowed us to target IFNαR1-deficient (LC^ΔIFNαR1^) or –sufficient LCs. These mice were transferred with TEα cells and then immunized with 1 μg of anti-huLangerin-hIgG4-Eα the next day. Ag-specific TEα cell responses were assessed 4 days later by flow cytometry. At fourteen days, flow cytometry and ELISA, respectively, assessed Ag-specific T cell and B cell responses. (**B**) Representative contour plots of TEα cells with the percentage of Tfh cells (PD1^+^CXCR5^+^) and summary graphs, including Bcl-6 MFIs of the proliferated TEα cells. (**C**) Representative contour plots of Ag-specific B cells with percentage of GC B cells (CD38^-^GL7^+^) and summary graph. (**D**) hIgG4-specific mIgG serum levels defined by ELISA. Data from two independent experiments were pooled. Each dot represents a separate mouse. (**E**) The treatment plan of Batf3^−/−^ mice with anti-IFNαR1 or isotype control antibody before and following LC targeting. (**F**) Left: the percentage of Tfh cells (PD1^+^CXCR5^+^) among proliferated TEα cells. Right: MFI of Bcl-6 of proliferated TEα cells. (**G**) Left: summary graph of Ag-specific GC B cells (CD38^-^GL7^+^), and right: hIgG4-specific mIgG serum levels defined by ELISA. Data from one representative experiment out of two is shown. Each dot represents a separate mouse. *p<0.05, ns=not significant.

To increase the rigor of our findings, we used a blocking antibody against IFNαR (**Fig. 2 E** and **Fig. S1 B**), which can address potential type-I interferon involvement in LC-induced adaptive immune responses in steady-state by acting directly on the T cells or indirectly through other cells. Again, we found no significant changes in T and B cell responses and a moderate increase rather than decrease or absence of anti-hIgG antibody levels (**Fig. 2, F and G**). Thus, these data strongly support that the adaptive immune responses induced by steady-state LCs are largely independent of type-I interferon signaling.

Next, we tested whether exogenous IFNα could boost adaptive immune responses above the levels induced by steady-state LCs. For this, we used an anti-muLangerin-IFNα construct we previously described that enables cDC1 to support antibody responses (Bouteau et al., 2019). LCs, similarly to cDC1, express IFNαR (**Fig. S1 A**). We found that delivering IFNα to LCs did not significantly alter their ability to support B cell responses (**Fig. S1 C**). In total, these data show that either promulgating or diminishing IFNα signaling in LCs does not affect their ability to support adaptive immune responses in this targeting model.

### IL-6 is not required for the induction of adaptive immune responses by steady-state LCs

IL-6 in mice, according to some but not all reports, plays a critical role in supporting the differentiation of Tfh cells in inflammatory models (Crotty, 2019). However, its role in Tfh cell and antibody response induction in a steady-state is unknown. Steady-state LCs, unlike cDC1s, contain high levels of IL-6 mRNA transcript (Bouteau et al., 2019). Since LCs, but not cDC1s, can induce Tfh cells with germinal center (GC) phenotype (Bcl-6^high^) and protective antibody responses in steady-state models (Yao et al., 2015; Bouteau et al., 2019), we hypothesized that LC-derived IL-6 might be essential in supporting the adaptive immune responses. To test this and limit IL-6 deficiency to LCs, we bred the IL-6^f/f^ mice (Sanchis et al., 2020) to the huLangerinCre mice (Kaplan et al., 2007) (**Fig. 3 A**). The resulting genotypes were determined using PCR, and the selective genomic recombination of the IL-6 locus in LCs was confirmed using PCR on sorted cells (**Fig. S1 D**). We then tested whether LC-derived IL-6 is needed to induce Tfh cells and antibody responses. The mice were transferred with transgenic CD4^+^ TEα cells and then injected with 1 μg of anti-huLangerin-Eα intraperitoneally, as described above. Cre-negative IL-6^f/f^ littermate mice served as controls. The mice were sacrificed 4 and 14 days later to characterize antigen-specific CD4^+^ T cell and B cell responses as presented above (**Fig. 3 A**). The expansion and phenotype of TEα cells in the absence of LC-derived IL-6 remained unchanged (**Fig. 3 B**). The B cell responses, including GC cells and anti-hIgG IgG serum levels, showed no significant changes (**Fig. 3 C**). Thus, these data support the idea that LC-derived IL-6 is not required to support steady-state adaptive immune responses.

**Figure 3.**
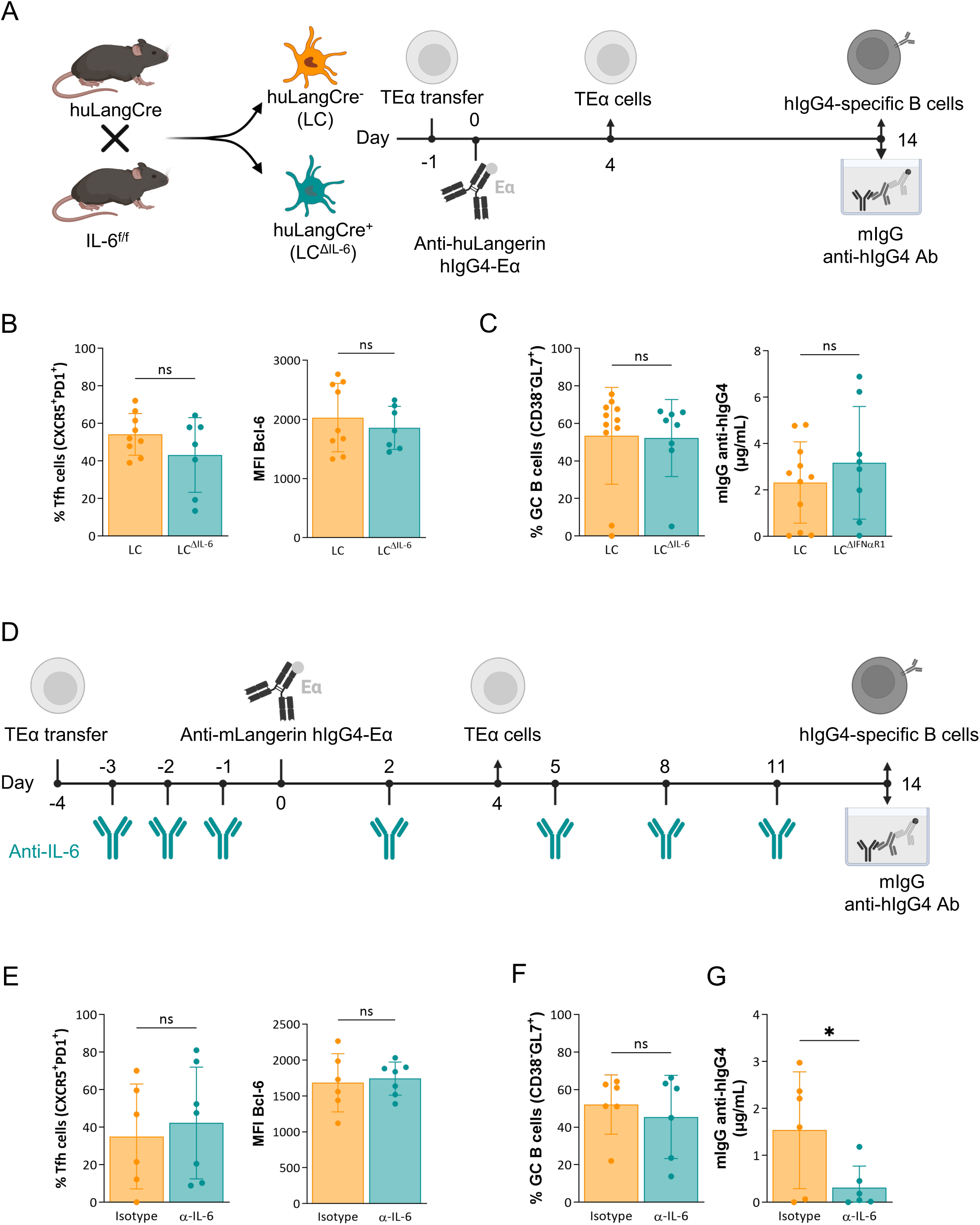
**IL-6 is not required for the induction of adaptive immune responses by steady-state LCs**. (**A**) huLangCre-IL-6^f/f^ mice were generated as depicted, which allowed targeting either IL-6-deficient (LC^ΔIL-6^) or IL-6-sufficient LCs. These mice were transferred with TEα cells and then immunized with 1 μg of anti-huLangerin-hIgG4-Eα the next day. Ag-specific TEα cell responses were assessed 4 days later by flow cytometry. At fourteen days, flow cytometry and ELISA assessed Ag-specific B cell responses. (**B**) Summary graphs on the percentage of Tfh cells (PD1^+^CXCR5^+^) and Bcl-6 MFIs of the proliferated TEα cells. (**C**) Summary graph on Ag-specific GC B cell percentages (CD38^-^GL7^+^), and hIgG4-specific mIgG serum levels defined by ELISA. Data from two independent experiments were pooled. Each dot represents a separate mouse. (**D**) The treatment plan of Batf3^−/−^ mice with anti-IL-6 before and following LC targeting. (**E**) Left: the percentage of Tfh cells (PD1^+^CXCR5^+^) among proliferated TEα cells. Right: MFI of Bcl-6 of proliferated TEα cells. (**F**) Summary graph of Ag-specific GC B cells (CD38^-^GL7^+^) and (**G**) hIgG4-specific mIgG serum levels defined by ELISA. Data from one representative experiment out of two is shown. Each dot represents a separate mouse. *p<0.05, ns=not significant.

To rule out the possibility that IL-6 produced and secreted by bystander cells might aid the induction of adaptive immune responses by steady-state IL-6 deficient LCs, we performed IL-6 blocking experiments. Batf3^−/−^ mice were treated with anti-IL-6 or isotype antibodies throughout the experiment (**Fig. 3 D**). Luminex^®^ assay on serum samples was used to confirm the efficiency of IL-6 blockade (**Fig. S1 E**). The antibody-treated Batf3^−/−^ mice that lack cDC1s were adoptively transferred with CD4^+^ TEα cells and then injected with 1 μg of anti-muLangerin-Eα, and then T and B cell responses were assessed as presented above. We found no significant differences in antigen-specific CD4^+^ T cell and B cell expansions and phenotype (**Fig. 3, D and F**). The anti-huIgG IgG ELISA on serum samples revealed only a moderate decrease in antibody levels in the anti-IL-6 treated mice (**Fig. 3 G**). Thus, cumulatively, our experiments showed that neither LC-derived nor total IL-6 plays a critical role in the induction of adaptive immune responses by steady-state LCs.

### ICOS-ICOS-L signaling has no major role in the induction of adaptive immune responses by steady-state LCs

Since steady-state levels of IL-6 and type-I interferon had no significant role in inducing adaptive immune responses by LCs, we turned our attention to membrane-bound co-stimulation. ICOS-ICOS-L signaling, including on DCs, is required to induce Tfh cells and subsequent antibody responses in inflammatory settings (Choi et al., 2011; Weber et al., 2015; Pratama et al., 2015). To determine whether ICOS-ICOS-L interaction is involved in the induction of Tfh cells and antibody responses by steady-state LCs, we exposed Batf3^−/−^ mice to ICOS-L blocking or isotype control antibodies (**Fig. 4 A, and S1 F**). The antigen-specific CD4^+^ T cell and B cell responses were characterized as discussed above. We found that blocking ICOS signaling minimally reduced the GC B cell but not antibody responses and did not affect the induction of GC-Tfh cells by steady-state LCs (**Fig. 4, B and C**). Thus, these data indicate that ICOS signaling is not critical in inducing adaptive immune responses by steady-state LCs.

**Figure 4.**
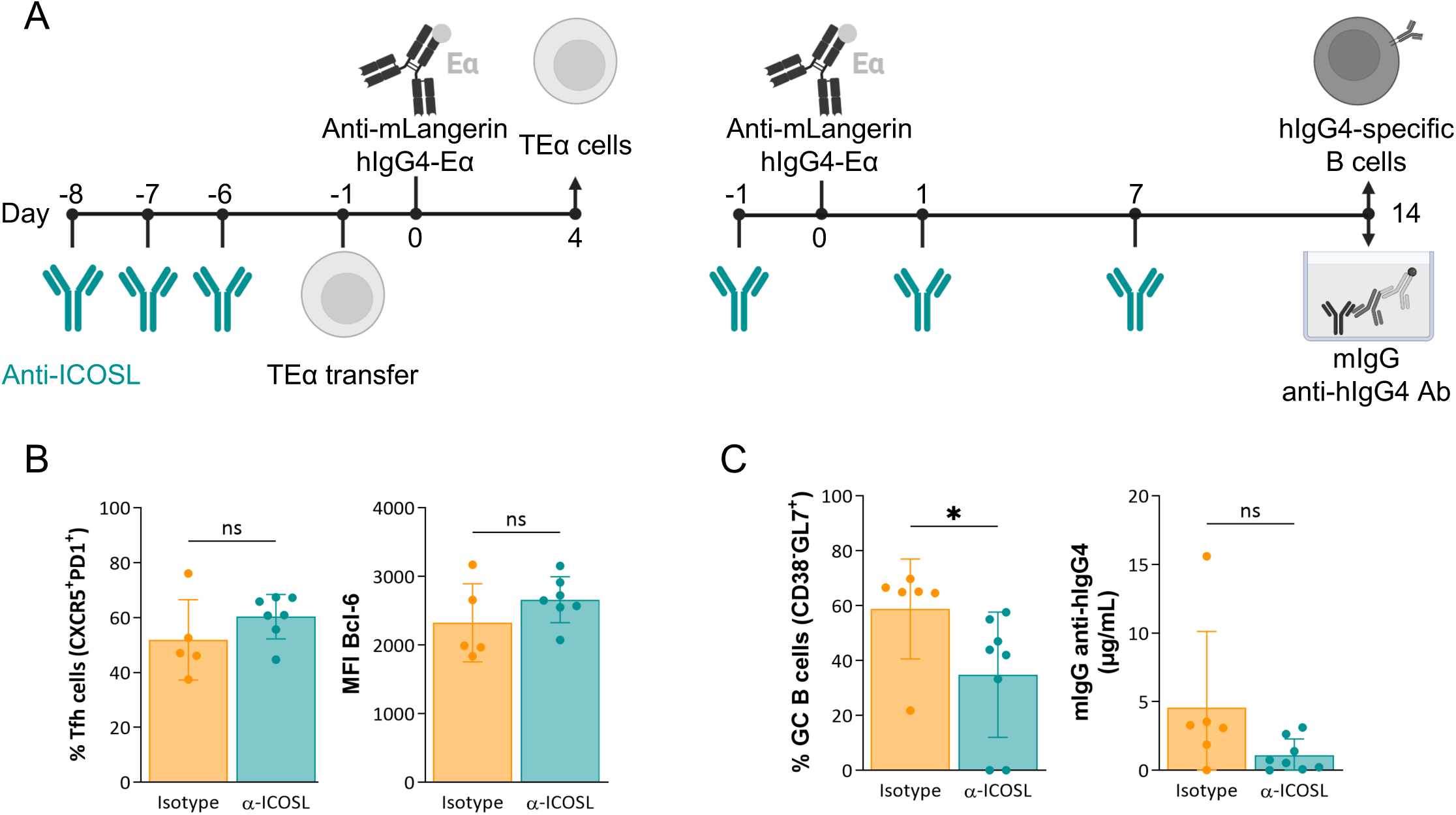
ICOS-ICOS-L signaling has no major role in the induction of adaptive immune responses by steady-state LCs. (**A**) The treatment plan of Batf3^−/−^ mice with anti-ICOS-L before and following LC targeting to assess T-(left) and B cell responses (right). (**B**) Left: the percentage of Tfh cells (PD1^+^CXCR5^+^) among proliferated TEα cells. Right: MFI of Bcl-6 of proliferated TEα cells. Data from one representative experiment out of three is shown. (**C**) Left: summary graph of Ag-specific GC B cells (CD38^-^GL7^+^), and right: hIgG4-specific mIgG serum levels defined by ELISA. Data from two independent experiments were pooled. Each dot represents a separate mouse. *p<0.05, ns=not significant.

### CD80/CD86 on DCs are critical for the induction of adaptive immune responses by steady-state DCs

Since the inflammatory cytokines and co-stimulation previously identified as critical for Tfh differentiation and antibody responses had no significant role in inducing adaptive immune responses by steady-state LCs, we decided to establish an *in vitro* steady-state model to more efficiently test for DC-derived factors involved in the induction of the adaptive responses. For this *in vitro* platform, we used the MutuDC1 DC cells line (Fuertes Marraco et al., 2012), OT-II cells from a Rag2^−/−^ background, and polyclonal primary B cells isolated from WT naïve mice (**Fig. 5 A**). The DCs and B cells were pulsed with OVA-peptide before co-culture, while cells not exposed to OVA served as controls. We anticipated that pulsing the polyclonal B cells with OVA-peptide should allow the cognate interaction with OT-II cells. Through this interaction, we expected that the B cells would provide the final maturation signals for the OT-II cells to differentiate into Tfh cells. In return, the T cells would facilitate the B cell responses, including isotype switching and antibody production. Indeed, we found that in the co-cultures where the DCs and the B cells were pulsed with OVA-peptide, the OT-II cells efficiently proliferated and differentiated into Tfh cells (**Fig. 5 B**). We also observed that a significant proportion of B cells underwent isotype switching and acquired GC phenotype (**Fig. 5 C**). We also detected substantial amounts of secreted IgG in the supernatant (data not shown). As expected, the inclusion of blocking MHC-II antibodies in the co-cultures prevented the T and B cell responses (**Fig. S2 A and B**). Furthermore, the primary DCs and MutuDC2 DC cell line (Pigni et al., 2018) in this co-culture assay also supported T and B cell responses but with different efficiencies (data not shown). Thus, we successfully established a steady-state co-culture system to study the induction of Tfh and B cell responses by DCs *in vitro*.

**Figure 5.**
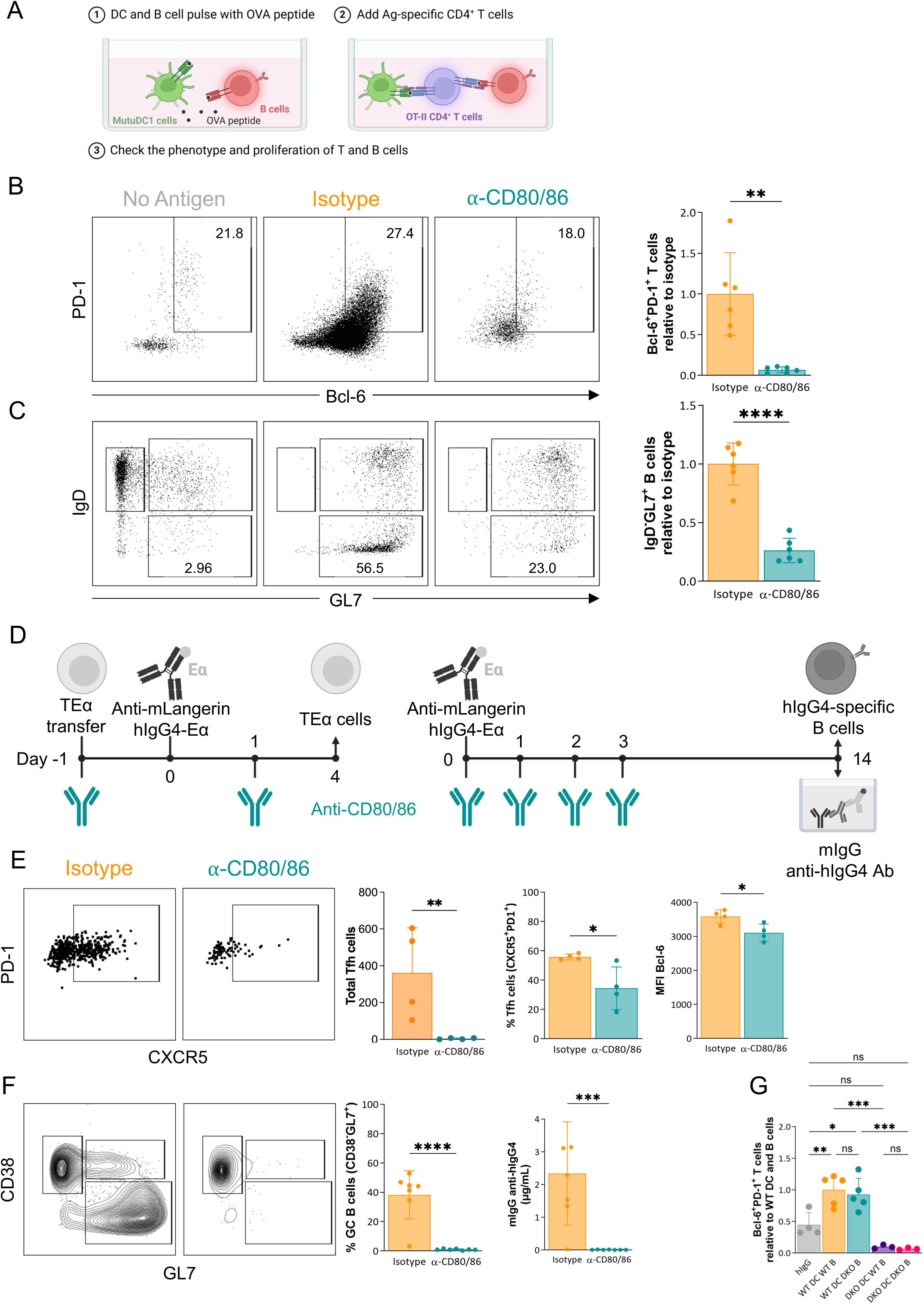
CD80/CD86 on DCs are critical for the induction of adaptive immune responses by steady-state DCs. (**A**) *In vitro* steady-state platform to model GC-dependent adaptive immune responses. (1) Murine DC cells (MutuDC1 cell line) and B cells enriched from WT mice were pulsed with OVA peptide. (2) Ag-specific CD4^+^ T cells (OT-II) were added after washing off the peptide. (3) After 5 days, the phenotype of T and B cells was determined by flow cytometry. (**B**) The role of CD80/86 in adaptive immune responses was tested *in vitro*. Anti-CD80/86 blocking Ab (or an isotype control Ab) were added simultaneously with T cells. Five days later, the phenotype of T cells was assessed by flow cytometry. Representative dot plot of proliferated OT-II cells with percentage of Tfh cells (PD1^+^Bcl-6^+^). Summary graph: the number of Tfh cells in each well was calculated and plotted relative to the average of Tfh cells in isotype conditions. Data from two independent experiments were pooled. Each dot represents a separate replicate. (**C**) Representative dot plot of proliferated B cells with percentage of GC cells (IgD^-^GL7^+^). Summary graph: the number of GC B cells in each well was calculated and plotted relative to the average of GC B cells in isotype conditions. Data from two independent experiments were pooled. Each dot represents an independent replicate. (**D**) The treatment plan of Batf3^−/−^ mice with anti-CD80/CD86 before and following LC targeting to assess T-(left) and B cell responses (right). (**E**) Representative TEα flow plots and summary graphs. Left: Total Tfh cells. Middle: The percentage of Tfh cells (PD1^+^CXCR5^+^) among proliferated TEα cells. Right: MFI of Bcl-6 of proliferated TEα cells. Data from one representative experiment out of three is shown. Each dot represents a separate mouse. (**F**) Representative flow plots and summary graphs for antigen-specific B cells. Left: summary graph of Ag-specific GC B cells (CD38^-^GL7^+^), and right: hIgG4-specific mIgG serum levels defined by ELISA. Data from two independent experiments were pooled. Each dot represents a separate mouse. (**G**) To test the role of CD80/86 on DCs and B cells, DCs and B cells were isolated from WT of CD80/86 DKO mice. DCs and B cells were pulsed for 24h with OVA peptide attached to an anti-mLangerin Ab (or non-targeted Ab (hIgG)). Then, after washing, Ag-specific OT-II T cells were added. After 5 days, the phenotype of T cells was checked by flow cytometry. The number of Tfh cells in each well was calculated and plotted relative to the average Tfh cells in WT DC/WT B cell conditions. Data from two independent experiments were pooled. Each dot represents an independent replicate. *p<0.05, **p<0.005, ***p<0.001, ****p<0.0001, ns=not significant.

Having established the steady-state co-culture system, we next tested whether membrane-bound costimulatory molecules are involved in the T and B cell responses. We supplemented the co-cultures with blocking antibodies targeting CD40L (CD154), CD80/CD86, and ICOS-L (CD275) to test. We observed that blocking CD80 and CD86, but not the others, significantly inhibited the differentiation of Tfh cells and class-switched (GL7^+^IgD^-^) B cell responses (**Fig. 5, B and C, and Fig. S2 A and B**). Thus, CD80 and CD86 are critical in adaptive responses induced by steady-state DCs *in vitro*.

To confirm that CD80 and CD86 are also crucial in LC-induced responses *in vivo*, we repeated our LC-targeting experiments presented above in the presence of anti-CD80/CD86 or corresponding isotype antibodies (**Fig. 5 D**). We found that antibodies blocking CD80/CD86 signaling *in vivo* significantly inhibited the induction of Tfh cells by LCs (**Fig. 5 E**) and entirely suppressed B cell responses, including GC formation and antibody production (**Fig. 5 F**). Thus, membrane-bound co-stimulation through CD80/CD86 is critical for steady-state LC-induced adaptive immune responses.

In inflammatory settings, CD80/CD86 on DCs, but not on B cells, are required for adaptive T and B cell responses (Watanabe et al., 2017). To define whether those findings also apply to our steady-state antigen targeting model, we first used our *in vitro* co-culture assay presented above with slight modification. We co-cultured magnetically enriched migratory DCs from the SDLNs of WT or CD80/CD86 double knock-out (DKO) mice pulsed with anti-muLangerin-OVA or IgG4-OVA with OT-II T cells and WT or CD80/CD86 double knock-out B cells (**Fig. S2 C and D**). The Tfh cell differentiation was unaffected by the lack of CD80/CD86 on the B cells but was almost absent in cultures with DCs that lacked CD80/CD86 (**Fig. 5 G**). These data, therefore, support the idea that CD80/CD86 expression by DCs plays a critical role in the induction of Tfh cell responses.

In summary, we demonstrate that Langerhans cells (LCs) promote GC-Tfh and B cell responses in steady-state, independent of IL-6, type-I interferon, and ICOS signaling. However, we found that CD80/CD86 expression on DCs is essential for inducing these adaptive immune responses. These findings differentiate steady-state T cell differentiation from the conventional three-signal model of T cell differentiation in inflammatory settings, where cytokines are indispensable third signals.

The induction of humoral immune responses in the steady-state is not exclusive to LCs. Splenic cDC1s have also been reported to support Tfh and antibody responses, though their ability to do so, unlike LCs’, depends on the receptor targeted (Steiner et al., 2022). In previous experiments, targeting migratory skin cDC1s through Langerin and Dectin-1 failed to elicit GC-Tfh or B cell responses, instead promoting cells with a pre-Tfh/Th1 phenotype (Bouteau et al., 2019). The reasons for these differences—whether due to distinct cDC1 subsets residing in different tissues (e.g., lymph nodes vs. spleen) or specific receptor targeting—remain to be elucidated. It will be crucial to investigate the instances when the cDC1s can support antibody responses, whether they undergo receptor-dependent alterations akin to those induced by inflammatory cues, such as IFN-α or poly-IC (Bouteau et al., 2019). As previously observed, cDC1-induced pre-Tfh/Th1 responses via Langerin targeting in steady-state were also independent of type-I interferons and IL-6 (Yao et al., 2015), further reinforcing that adaptive immune response induced by DCs in steady-state are independent of cytokines. Notably, in our model, no polarizing adjuvants or pattern recognition ligands were used that could directly or indirectly differentially affect LCs and cDC1s. Both subsets were targeted via the same receptor, yet they induced distinct adaptive immune responses. This strongly supports the idea that DC subsets are functionally specialized or pre-programmed, even in steady-state, to drive specific adaptive immune responses. Our findings align with studies on pre-committed DC precursors in the bone marrow (Satpathy et al., 2012; Schlitzer et al., 2015) and human *in vitro* observations that both primary tissue– and CD34+ cord-blood-derived LCs, but not monocyte-derived DCs support Tfh and B cell responses in steady-state (Bouteau et al., 2019; Kervevan et al., 2021), suggesting that tissue residency might play a limited role in shaping the functional specialization of DC subsets. Also, these results underscore the intrinsic programming of DC subsets in directing distinct immune pathways and, at the same time, highlight the translatability of mouse data to humans.

The findings that these prominent immunological factors in defining Tfh cell differentiation and B cell responses in our model do not seem to be critical contributors do not rule out that other cytokines or soluble factors might play a significant role in the process. However, our observations may offer insight into the mechanisms underlying homeostatic immunity toward commensals (Belkaid and Harrison, 2017) and the development of certain autoimmune diseases. Engagement of DCs by commensals through pattern recognition receptors, unlike pathogenic interactions, typically leads to no detectable or minimal activation and maturation (Ansaldo et al., 2021), suggesting limited involvement of DC-derived cytokines in polarizing commensal-specific T helper (Th) subsets. If cytokines are not critical in steady-state, what drives distinct Th responses? One possibility is that DC subsets differ in their peptide-MHC levels, a hypothesis aligned with the quantitative model proposed by van Panhuys and colleagues (Van Panhuys, 2016). However, our findings suggest that additional factors beyond peptide-MHC levels contribute to Tfh and B cell responses. LCs, across a wide range of antigen doses, while with different efficiency, uniquely support antibody responses (Yao et al., 2015; Bouteau et al., 2019), unlike cDC1s, which fail to do so under similar conditions. LCs appear to provide stronger cumulative TCR stimulation, evidenced by sustained CD69 expression, pS6 phosphorylation (Bouteau et al., 2019), and Nur77-GFP signals (unpublished observation) in T cells activated by LCs. Given that Tfh differentiation requires stronger TCR stimulation than Th1 differentiation (Bhattacharyya and Feng, 2020), this enhanced signal may explain the unique ability of LCs to promote Tfh cells. How LCs provide more potent stimuli than cDC1s and drive distinct adaptive immune responses in steady-state and inflammation (Igyártó et al., 2011) remains to be determined. While here we show that CD80/CD86 play an essential role in LC-induced adaptive immune responses, the contribution of CD80/CD86 as the second signal is likely not unique to LCs since other DCs in the lymph nodes also express high levels of CD80/CD86. Thus, other factors unique to these DC subsets, such as differentially expressed CD11a and CD11b integrins (Bouteau et al., 2019), molecules that play an essential role in regulating immunological synapses (Balkow et al., 2010; Varga et al., 2007), could likely serve as the decisive “third signal” and polarize the naïve T cells into distinct Th cells.

LCs and cDC1s exhibit distinct cytokine transcript profiles in the steady state, which may underlie their functional specialization (Igyártó et al., 2011). LCs are enriched in transcripts for *Il1b, Il6*, and *Il23p19*, aligning with their capacity to promote Th17 responses. Conversely, cDC1s harbor higher levels of *Il12p40* and *Il27* transcripts, correlating with their roles in driving Th1 and CTL responses. Interestingly, *Il12p40* transcript expression is detectable even in pre-committed bone marrow cDC1 precursors (Schlitzer et al., 2015), indicating intrinsic programming that primes DCs for rapid and specific cytokine production upon activation. These findings suggest that DC subsets are poised to respond swiftly to external stimuli by producing subset-specific cytokines that facilitate targeted Th polarization. However, our steady-state results indicate that cytokines may not be involved in the polarization of Th cells under non-inflammatory conditions. Therefore, we propose a model wherein, in the steady-state differential expression of adhesion molecules—such as integrins (CD11a and CD11b)—by DC subsets may provide a critical “third signal” to skew adaptive immune responses. These integrin-mediated could selectively promote “basic” Tfh, Th1, and potentially Th2 differentiation without the need for cytokine-mediated signaling. Under inflammatory conditions, cytokines would act as a “fourth signal,” boosting and further polarizing the T helper cells into effector subsets, such us Th17, Th22, and others, tailored to combat the specific infection and restore tissue homeostasis. This dual-layered model highlights how DC subsets integrate their intrinsic programming with environmental cues to orchestrate adaptive immunity dynamically.

## FIGURE LEGENDS

**Figure S1**. **Validation experiments for type I interferon, IL-6, and ICOS-L interference.** (**A**) huLangCre-IFNαR1^f/f^ mice were generated to delete IFNαR1 in LCs specifically. SDLN of huLangCre-IFNαR1^f/f^ mice (Cre+), littermate controls (Cre-) and IFNαR1 complete KO mice were stained for IFNαR1 by flow cytometry. MFI of IFNαR1 was calculated for B cells, cDC1s, LCs, and rDCs. Data from multiple experiments pooled together. Each dot represents a separate mouse. (**B**) Batf3^−/−^ mice were treated with anti-IFNαR1 blocking Ab or an isotype. Four and fourteen days after LC targeting, SDLN were isolated and stained for IFNαR1 (same clone used *in vivo* to block the receptor). Left: representative histogram of B cells from isotype or anti-IFNαR1 treated mice. Shaded grey are B cells stained with an isotype control. Right: summary data of IFNαR1 staining on B cells. (**C**) LCs were targeted with 1 μg of anti-mLangerin-hIgG4 Ab in the absence or presence of IFNα. Fourteen days later, the percentage of GC-B cells among hIgG4 specific B cells was assessed by flow cytometry (left), and anti-hIgG4 Ab responses (right) were assessed by ELISA on serum. Data from two experiments pooled. Each dot represents a separate mouse. (**D**) huLangCRE-IL-6^f/f^ mice were generated to target IL-6-deficient (LC^ΔIL-6^) or –sufficient LCs. LCs and keratinocytes (KC) of Cre+ and Cre-mice were sorted, and genomic DNA was extracted for genotyping. Note that the recombined band is only present in Cre+ LCs. (**E**) The serum of Batf3^−/−^ mice treated with anti-IL-6 or isotype control antibodies was collected, and the concentration of IL-6 was assessed by Luminex. (**F**) The efficiency of ICOS-L blockade is shown. B cells were stained with anti-ICOS-L. Each dot represents a separate mouse. **p<0.005, ***p<0.001, ****p<0.0001, ns=not significant.

**Figure S2**. **Validation experiments for CD80/CD86 and other co-stimulatory molecules.** (**A**) The role of MHCII, CD154, CD275, CD80/86, or a combination of these parameters in GC responses was tested *in vitro*. Blocking Abs or isotype control Abs were added to the *in vitro* model simultaneously with T cells. Five days later, the phenotype of T cells (left) and B cells (right) was assessed by flow cytometry. The Tfh and GC B cells in each well were calculated and plotted relative to the average of Tfh and GC B cells in isotype conditions. Data from two independent experiments were pooled. Each dot represents an independent replicate. (**C**) At the end of the in vitro cultures with WT and CD80/86 DKO DCs and B cells, the level of CD86 on DCs and (**D**) B cells was determined by flow cytometry, and the MFI values of CD86 were plotted. Data from two independent experiments were pooled.

## MATERIALS AND METHODS

### Mice

huLangerin (also called huLangerin-DTR) (Bobr et al., 2010), Batf3^−/−^ (Edelson et al., 2010), and huLangerinCre (Kaplan et al., 2007) mice have been previously described. huLangerinCre were crossed with IFNαR1^f/f^ (Jackson Laboratories strain 028256) or IL-6^f/f^ (mice provided by Dr. Roger Davis, University of Massachusetts; developed by Dr. Juan Hidalgo at Universitat Autònoma de Barcelona) (Sanchis et al., 2020). CD90.1 congenic TEα Rag1^−/−^ CD4 TCR transgenic mice to I-Eα_52−68_ on the C57BL/6 background were initially obtained from Dr. Marc Jenkins (University of Minnesota). IRF8^32Δ^ mice were obtained from Dr. Kenneth Murphy (Washington University School of Medicine) and Jackson Laboratories. XCR1-Cre-DTA mice were generated by crossing XCR1-Cre (Jackson Laboratories strain 035435; developed by Dr. Kenneth Murphy) with “STOP”-DTA (Jackson Laboratories strain 009669). WT C57BL/6J (strain 000664), CD45.1 PepBoy (strain 002014), and OT-II (strain 005194) were purchased from Jackson Laboratories and maintained in our facility. Rag2^−/−^ OT-II mice were purchased from Taconic model 11490. CD80/86 double knock-out (DKO) mice developed by Dr. Arlene H. Sharpe (Mass General Hospital and Harvard Medical School) (Borriello et al., 1997) was provided by Drs. Masashi Watanabe and Richard Hodes (NIH). All experiments were performed with 6-to 12-week-old female and male mice. Mice were housed in microisolator cages and fed autoclaved food and water. The Institutional Care and Use Committee at Thomas Jefferson University approved all mouse protocols under protocol number: 02315.

### Steady-state Langerin targeting

These experiments were performed with anti-human/anti-mouse mAb and conjugates (anti-huLangerin-Eα, anti-muLangerin-Eα anti-muLangerin-doc, cohesin-IFNα4) generated in-house, as previously described (Bouteau et al., 2019). All the reagents used in this study were generated using mammalian cell lines to minimize the presence of endotoxins. The average endotoxin level was below 0.2 ng LPS/mg protein. Eα (I-Eα_52−68_) is a well-characterized immunodominant T cell epitope from the I-Eα MHCII molecule recognized by transgenic TEα cells in the context of I-A_β_. For the generation of the anti-muLangerin-IFNα4 construct, we used a previously described technology that relies on the high-affinity interactions between dockerin (doc) and cohesin (coh) (Bouteau et al., 2019). Dockerin was fused to the heavy chain of the antibody. Mice received intravenous (i.v.) transfer of CFSE-labeled, congenically marked 3×10^5^ TEα cells 1 day before antigen targeting as previously described (Bouteau et al., 2019). One μg of anti-human/anti-mouse mAb and conjugates or PBS were administered intraperitoneally (i.p.) on day 0. For TEα cells characterization, mice were sacrificed 4 days after Langerin targeting, and the skin-draining lymph nodes (SDLNs; axillary, brachial, and inguinal) were harvested for flow cytometry. For B cell characterization, mice were sacrificed on day fourteen, the SDLNs were harvested for flow cytometry, and the blood was collected for serum isolation and ELISA. For dendritic cell characterization, SDLNs were digested as described previously (Bouteau et al., 2019) before staining for flow cytometry.

### Flow cytometry

Flow cytometry was performed on single-cell suspensions of the SDLNs (axillary, brachial, and inguinal) of mice. The fixable viability dye (Thermo Fisher Scientific) was used to exclude dead cells. The following antibodies from BioLegend were used to stain cells CD3 (145-2C11), CD4 (GK1.5), CD16/CD32 (93), CD19 (6D5), CD25 (PC61), CD44 (IM7), CD45.2 (104), CD69 (H1.2F3), CD80 (16-10A1), CD86 (GL-1), CD90.2 (30-H12), CD138 (281-2), CD275 (HK5.3), B220 (RA3-6B2), Blimp1 (5E7), CXCR5 (L138D7), GL7 (GL7), IFNaR1 (MAR1-5A3), IgD (11-26c.2a), IgM (RMM-1), MHCII (M5/114.15.2), PD-1 (29F.1A12),and Sca1 (D7). CD4 (GK1.5), CD38 (90), CD90.1 (OX-7), Bcl6 (K112-91), and IgD (D7) were purchased from BD Biosciences. CD11b (M1/70), CD90.2 (53-2.1), F4/80 (BM8) were purchased from Thermo Fisher Scientific. For antigen-specific B cell staining, cells were also incubated *ex vivo* with AF647-conjugated targeting Ab construct. Intracellular transcription factor staining was performed with the BD Bioscience Cytofix/Cytoperm kit (BD Biosciences, San Jose, CA), according to the manufacturer’s instructions. All the flow cytometric plots presented in this article were pre-gated on live (using Live/Dead stain) and singlet events. Samples were analyzed on an LSRFortessa or Symphony flow cytometer (BD Biosciences), and data were analyzed with FlowJo software (BD Biosciences).

### Assessment of Humoral Immune Responses by ELISA

Serum samples were obtained 14 days after immunization with anti-Langerin Ab constructs using BD Microtainer SST tubes (BD) and stored at −80°C. To determine antigen-specific antibody titers, clear flat-bottom immune 96-well plates were coated with 50 μL of huIgG4 protein diluted in BupH Carbonate-Bicarbonate buffer (Thermo Fisher Scientific) at 2 μg of protein/ml and incubated overnight at 37C. After washing, plates were blocked with blocking buffer (TBS; Thermo Fisher Scientific). After blocking, the buffer was discarded, and serial dilutions of serum in the blocking buffer were added and incubated for 2 h at 37°C. A serial dilution of a mouse anti-hIgG4 antibody (EMD Millipore) was used as a standard. After washing, plates were incubated with horseradish peroxidase (HRP)–conjugated goat anti-mouse IgG (Jackson ImmunoResearch; West Grove, PA) in blocking solution for 2 h at 37°C, washed and developed with HRP substrate (Ultra-TMB Chromogen Solution: ThermoFisher Scientific). The reaction was stopped with 1N HCl and plates were read at 450 nm.

### IFNαR1 antibody *in vivo* treatment

For blockade of IFNαR1, Batf3^−/−^ mice were treated with 0.5 mg/mouse of IFNαR1 blocking antibody (clone MAR1-5A3; BioLegend) on 3 consecutive days pre-immunization and 0.25 mg every 3 days post-immunization (Macal et al., 2018). Control mice received a similar amount of a mouse IgG1 isotype control antibody (clone MOPC-21; BioLegend). All antibodies were administrated i.v. in 300 μl of PBS. To validate the block, SDLNs were isolated and stained for IFNαR1 (the same clone used *in vivo* to block the receptor).

### IL-6 antibody *in vivo* treatment

For the blockade of IL-6, Batf3^−/−^ mice were treated with 0.5 mg/mouse of IL-6 blocking antibody (clone MP5-20F3; BioLegend) on one-day pre-immunization and 0.25 mg every other day post-immunization. Control mice received a similar amount of a rat IgG1 isotype control antibody (clone RTK2071; BioLegend). All antibodies were administrated i.v. in 300 μl of PBS. To validate the block, the blood of each mouse was collected, and serum was isolated for Luminex^®^ analysis.

### ICOS-L antibody *in vivo* treatment

For blockade of ICOS-L (CD275) and TEα cell characterization, Batf3^−/−^ mice were treated with 0.1 mg/mouse of ICOS-L blocking antibody (clone HK5.3; BioLegend) on days –8, – 7, and –6 pre-immunization. CFSE-labeled TEα cells were transferred on day –1. Control mice received a similar amount of a rat IgG2a isotype control antibody (clone RTK2758; BioLegend). All antibodies were administrated i.v. in 300 μl of PBS. For blockade of ICOS-L (CD275) and B cell characterization, Batf3^−/−^ mice were treated with 0.1 mg/mouse of ICOSL blocking antibody (clone HK5.3; BioLegend) on day –1 pre-immunization, and days 1 and 7 post-immunization. Control mice received a similar amount of a rat IgG2a isotype control antibody (clone RTK2758; Biolegend). All antibodies were administrated i.v. in 300 μl of PBS. To validate the block, the blood of each mouse was collected and stained for ICOS-L with the exact clone as the antibody used for the *in vivo* treatment.

### CD80/86 antibodies *in vivo* treatment

For blockade of CD80 and CD86 and TEα cell characterization, Batf3^−/−^ mice were treated with 0.15 mg/mouse of CD80 (clone 16-10A1; BioXCell) and 0.15 mg/mouse of CD86 (clone GL-1; BioXCell) blocking antibodies on days –1, 0, and 1 (relative to immunization day). Control mice received a similar amount of a rat IgG2a isotype control antibody (clone RTK2758; BioLegend). All antibodies were administrated i.p. in 100 μl of PBS. For blockade of CD80 and CD86 and B cell characterization, Batf3^−/−^ mice were treated with 0.15 mg/mouse of CD80 (clone 16-10A1; BioXCell) and 0.15 mg/mouse of CD86 (clone GL-1; BioXCell) blocking antibodies on days 0, 1, 2 and 3. Control mice received rat IgG2a and Armenian hamster IgG isotype control antibodies (clones 2A3 and BE0091, respectively; BioXCell). All antibodies were administrated i.p. in 100 μl of PBS. SDLNs were stained with CD80 and CD86 antibodies to validate the block (same clone as the antibodies used for the *in vivo* treatment).

### *In vitro* assay with MutuDC1

For this assay, we combined and optimized protocols from P. Sage and A. Sharpe (Sage and Sharpe, 2015) and Kolenbrander et al., (Kolenbrander et al., 2018). Cells were prepared in co-culture in 96-well U-bottom plate as follows: 1) (**ref**) 10^4^ MutuDC1 cells/well were distributed in complete IMDM (IMDM w/ Glutamax w/ HEPES, 8% Heat-inactivated FBS, 50 μM Δ-mercaptoethanol and 1% pen/strep). 2) B cells were labeled with Cell-Trace Yellow (BioLegend) and enriched (Mojosort pan-B cell selection kit from BioLegend) from the spleen of CD45.1 mice. 2.5×10^5^ CD45.1 B cells were distributed in the same wells as MutuDC1. 3) MutuDC1 and B cells were pulsed at 15 μg/mL with OVA_323-339_ peptide (GenScript) for 1 hour at 37°C 5% CO_2_. 4) Rag2/OT-II T cells were isolated from spleen, mesenteric and SDLNs and labeled with Cell-Trace Violet (BioLegend). After washing MutuDC1 and B cells with complete IMDM twice, 5×10^5^ Rag2/OT-II T cells were added per well. 5) After mixing, cells were incubated for 5 days at 37°C 5% CO_2_.

Different controls lacking different cell populations or peptides were used. If any blocking antibodies were added to wells, it was at the same time as the addition of T cells at 5 μg/mL. Blocking antibodies used were MHC-II block (clone Y-3P from BioXCell), CD80 (clone 16-10A1 from BioLegend); CD86 (clone GL-1 from BioLegend); CD154 (clone MR1 from BioLegend); CD275 (clone HK5.3 from BioLegend). At the end of the incubation, the plate was centrifuged. The supernatant was used for ELISA mIg detection. Briefly, clear flat-bottom immune 96-well plates were coated with 50 μL of F(ab’)2 Donkey anti-mouse IgG (H+L) (ThermoFisher Scientific) diluted at 8 μg/mL in BupH Carbonate-Bicarbonate buffer (Thermo Fisher Scientific) and incubated overnight at 37°C. After washing with PBS + Tween20 (ThermoFisher Scientific), plates were blocked with a 2% milk solution for 30 minutes at 37°C. After washing, serial dilutions of supernatant in TBS were added and incubated for 2 h at 37°C. After washing, plates were incubated with horseradish peroxidase (HRP)–conjugated Donkey anti-mouse IgG (H+L) (ThermoFisher Scientific) in TBS for 2 h at 37°C, washed and developed with HRP substrate (Ultra-TMB Chromogen Solution: ThermoFisher Scientific). The reaction was stopped with 1N HCl and plates were read at 450 nm.

The cells were stained with the following antibodies: CD4 (GK1.5; BD Biosciences), CD44 (IM7; BioLegend), CD45.1 (A20; BioLegend), PD1 (39F.1A12; BioLegend), Bcl6 (K112-91; BD Biosciences), GL7 (GL7; BioLegend), IgD (11-26c.2a; BioLegend), CD19 (6D5; BioLegend), CD16/30 (93; BioLegend), and a fixable live/dead from Thermofischer Scientific.

### *In vitro* assay with primary DCs

This assay is similar to the *in vitro* assay with MutuDC1 with the following differences. The primary DC fraction was a CD11c positive enrichment (Mouse CD11c positive selection kit II from StemCell) of SDLNs of WT and CD80/86 DKO mice. The T cell fraction is a CD4 T cell enrichment (Mojosort mouse CD4 T cell enrichment from BioLegend) of spleen mesenteric and SDLN of OT-II mice. The B cell fraction is a B cell enrichment (Mouse Pan-B cell enrichment from BioLegend) of WT and CD80/86 DKO mice spleens. The antigen used here is anti-muLangerin-doc/cohesin-OTII (and control IgG4-doc/cohesion-OTII) generated in-house, as previously described (Bouteau et al., 2019). Primary DC (5×10^4^ cells) and B cell (10^6^ cells) fractions were pulsed for 24 hours at 37°C 5% CO_2_ with 10nM of the targeting or control constructs. After washing, 10^5^ cells of the T cell fraction were added and incubated for 5 more days at 37°C 5% CO_2_.

The staining of the cells at the end of the co-culture assay included the following antibodies from BD Biosciences: CD4 (GK1.5), CD38 (90) and Bcl-6 (K112-91). ThermoFisher Scientific antibodies used were specific for MHCII (M5/114.15.2), CD4 (GK1.5), CD86 (GL1), and the fixable live/dead marker. The following antibodies were purchased from BioLegend: CD44 (IM7), CD45.1 (A20), CD138 (281-2), IgD (11-26c.2a), CD19 (6D5), GL7 (GL7), CD11c (N418), PD1 (29F.1A12), and CD16/32 (93).

### Statistical Analysis

Differences between 2 data sets were analyzed first for normality using the Shapiro-Wilk test. If normality was met, an unpaired t-test was used to assess the difference between the 2 different data sets. In the absence of normality, a Mann-Whitney test was used. For data sets with more than two groups, an ordinary one-way ANOVA with multiple comparisons was used. All analysis were performed with GraphPad Prism software.

## ACKNOWLEDGMENTS

We thank the BioImaging Shared Resources, Flow Cytometry, and Laboratory Animal Facilities at Sidney Kimmel Cancer Center for their expert assistance in performing the presented experiments. We thank Dr. Hans Acha-Orbea for sharing the MutuDC cell lines. We thank Dr. Luis Sigal for sharing the IFNαR^KO^ mice, Drs. Arlene Sharpe, Masashi Watanabe, and Richard Hodes for the CD80/CD86^DKO^ mice, Drs. Juan Hidalgo and Roger Davis for the IL-6 floxed mice, and Dr. Kenneth Murphy for the IRF8^32Δ^ mice. Jerome Ellis and Zhiqing Wang are thanked for reagent making, and Jiukai Yu for laboratory assistance. Some of the figures were generated using BioRender. This work was supported by the National Institute of Allergy and Infectious Diseases (https://www.niaid.nih.gov) R01AI146420 to B.Z.I. and institutional start-up funds. The Flow Cytometry and Laboratory Animal Facilities at Sidney Kimmel Cancer Center at Thomas Jefferson University are supported by the National Cancer Institute (https://www.cancer.gov) P30CA056036.

## AUTHORS CONTRIBUTIONS

A.B., Z.Q., and B.Z.I. designed experiments. A.B. and Z.Q. performed experiments, analyzed the data, and generated the figures. S.Z. and G.Z. supervised in-house reagent making and quality assurance and performed Luminex assays. B.Z.I. wrote the manuscript. All authors participated in discussions of experimental results, edited the manuscript, and approved the final manuscript.

