## Supplementary figures and images for "Langerhans Cells Drive Tfh and B Cell Responses Independent of Canonical Cytokine Signals"

### Supplemental Figure 1

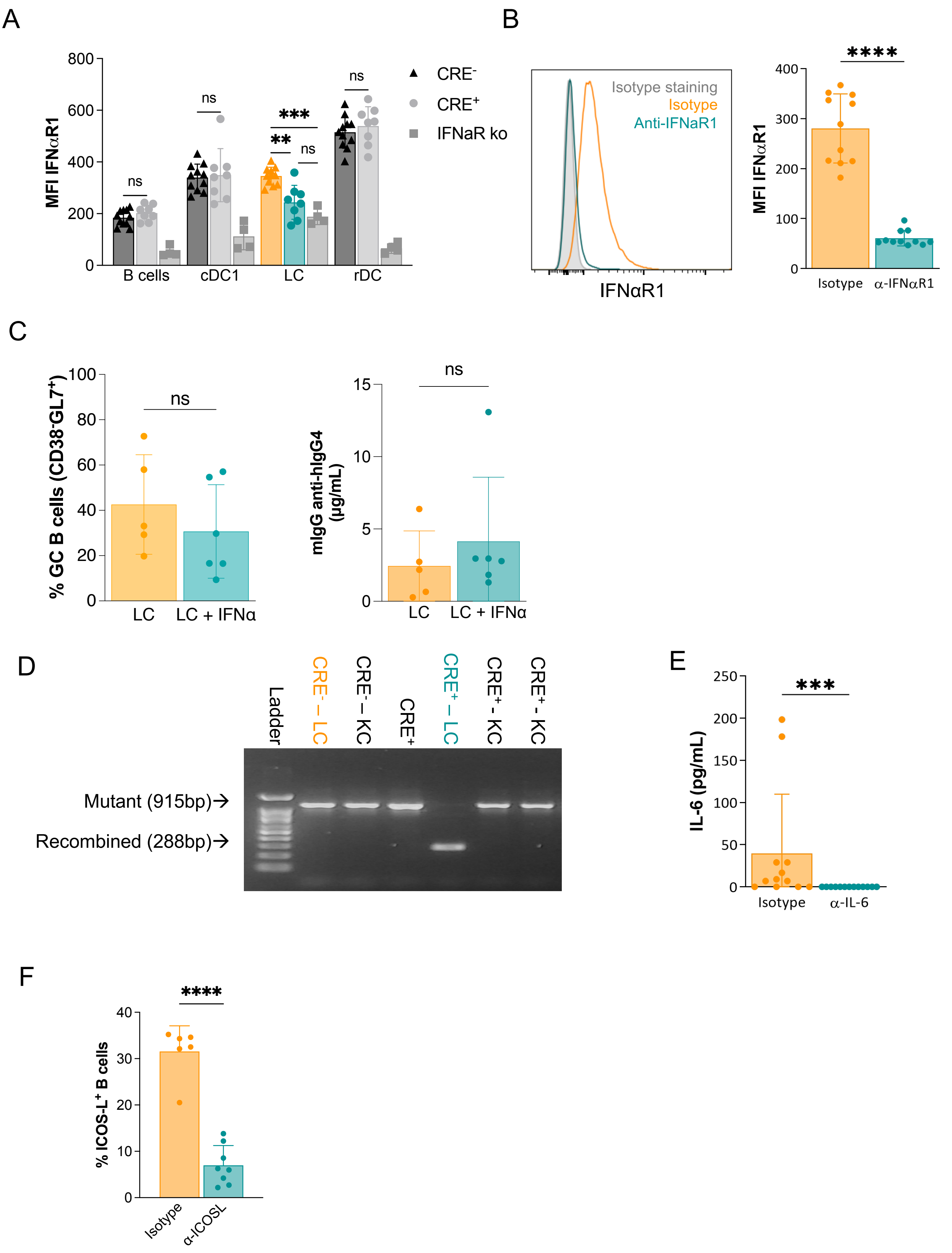

### Supplemental Figure 2

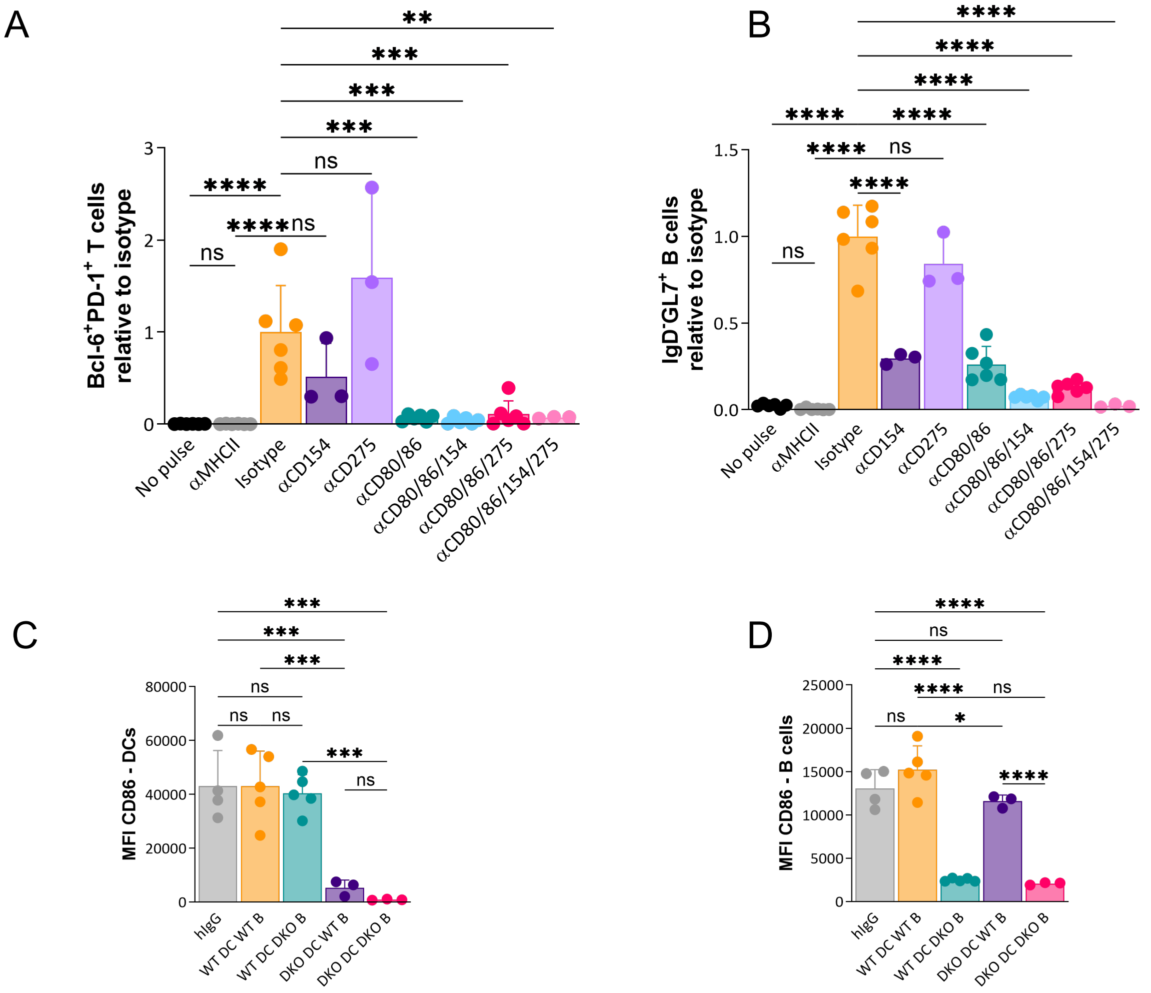
